# Viral rewiring of DDR signaling activates a pro-survival network that drives chemotherapy resistance

**DOI:** 10.64898/2026.07.06.736708

**Authors:** Michael A. Thomas, Jiang Kong, Hafiz Muhammad Faraz Azhar, Ahlam Akmel, Fayuan Wen, Jinrui Xu

## Abstract

Radiation and chemotherapy rely on an intact DNA damage response (DDR) to halt cell-cycle progression and eliminate damaged cells, yet many tumors evade these outcomes and develop resistance. Adenoviruses remodel host signaling networks in ways that mirror tumor evolution, providing a powerful system to dissect how DDR pathways are subverted. Here, we identify two scenarios in which the central DDR kinases ATM and ATR are reprogrammed from enforcing CHK1/CHK2-dependent checkpoint arrest to activating a NEMO-NF-κB survival pathway. This rewiring induces transcriptional programs associated with stress tolerance, anti-apoptotic signaling, and chemoresistance, and promotes the accumulation of cells with abnormal DNA content. These findings reveal a previously unrecognized mode of DDR plasticity that generates a pro-survival state reminiscent of early tumor evolution and suggest how ATM- and ATR-dependent pathways can be co-opted to promote therapeutic resistance.

## INTRODUCTION

The DNA damage response (DDR) is initiated when the MRN complex, composed of MRE11, RAD50, and NBS1, recruits the apical kinases ataxia-telangiectasia mutated (ATM) and ATM-and-Rad3-related (ATR) to sites of DNA lesions, triggering an evolutionarily conserved signaling network that preserves genome integrity [1–3]. Once activated, ATM and ATR phosphorylate key transducers, including the checkpoint kinases CHK1 and CHK2, and downstream effectors such as p53, thereby coordinating cell-cycle arrest, DNA repair, or, when damage is irreparable, cell death [1–3]. Cytotoxic therapies, including radiation and chemotherapy, exploit this signaling network to eliminate tumor cells, making intact DDR function essential for therapeutic efficacy. Yet many tumors evade these outcomes [4], underscoring persistent gaps in our understanding of how DDR pathways are subverted during therapy resistance.

Adenoviruses (Ads) are widespread human pathogens, whose replication cycle is well characterized. Although not oncogenic in humans, Ads remodel host signaling networks in ways that mirror early tumor evolution, making them powerful tools for dissecting how cancer-relevant pathways are rewired [5, 6]. Ad proteins perturb numerous cellular systems—including the DDR, NF-κB signaling, and cell-cycle control-that are frequently dysregulated in cancer [7, 8].

Here, we use adenovirus infection to uncover two scenarios in which the central DDR kinases ATM and ATR are reprogrammed from enforcing CHK1/2-dependent checkpoint arrest to activating a NEMO-NF-κB survival pathway. This signaling switch promotes the accumulation of cells with abnormal DNA content and induces transcriptional programs associated with stress tolerance, anti-apoptotic signaling, and chemoresistance. These findings reveal an unappreciated plasticity in DDR signaling that generates a pro-survival state reminiscent of early tumor evolution and suggest how ATM- and ATR-dependent pathways can be co-opted to promote therapeutic resistance and potentially limit the efficacy of Ad-based vaccines and oncolytic agents.

## RESULTS

### E4orf3 rewires ATM-ATR checkpoint signaling to drive abnormal DNA content

Adenovirus infection is widely assumed to inactivate ATM and ATR signaling because the virus disables upstream DNA damage sensing [9, 10]. The E1B55K-E4orf6 complex targets MRE11 and RAD50 for proteasomal degradation, while E4orf3 sequesters MRE11 and NBS1 into higher-order nuclear assemblies [11], collectively disrupting the MRN complex and preventing canonical DDR activation. Despite this, ATM phosphorylation at Ser1981-a hallmark of ATM activation [12]-has been consistently observed in Ad-infection [13–15], prompting us to examine how ATM and ATR behave when their upstream sensors are disabled.

Although HeLa cells harbor chromosomal translocations and other structural abnormalities not typically present in normal human cells [24], flow-cytometric analysis (Fig. 1A) showed expected G1, S, and G2/M distributions in mock-infected populations, with only ∼1% of cells exceeding 4N DNA content (DNA >4N). *E4orf3(+)* Ad Infection dramatically increased the fraction of cells with abnormal DNA content, whereas *E4orf3(-)* Ad induced a pronounced G2/M arrest (Fig. 1A, B). These contrasting phenotypes suggested that E4orf3 fundamentally alters genome-content control.

**Figure 1.**
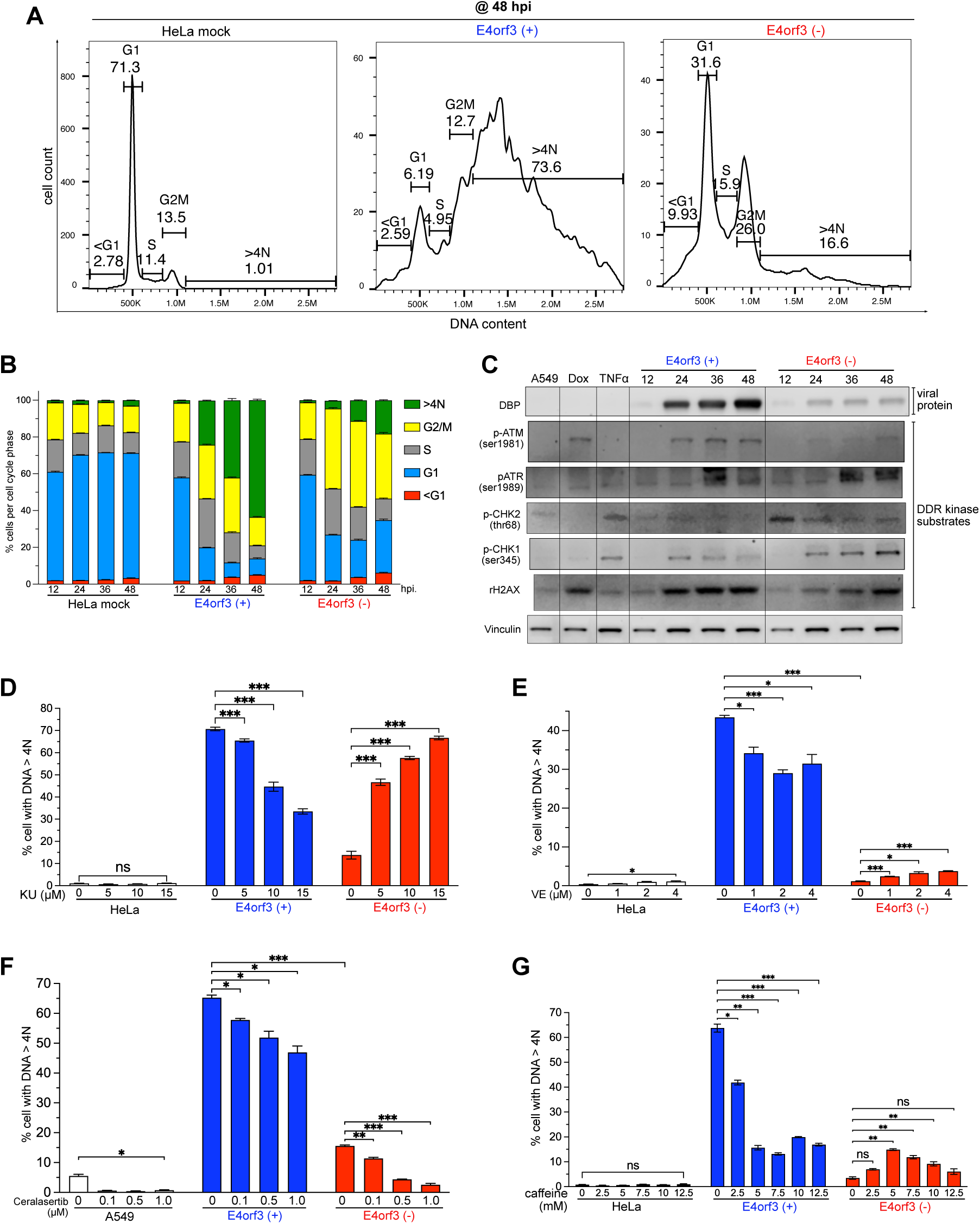
E4orf3 rewires ATM-ATR checkpoint signaling to drive abnormal DNA content. **(A)** Representative DNA-content histograms of HeLa cells mock-infected or infected with *E4orf3(+)*, or *E4orf3(-)* adenovirus (Ad) at 48 hpi. Cells with DNA > 4N indicate abnormal DNA content. **(B)** Time-course analysis of cell-cycle distributions in mock-infected, *E4orf3(+)* or *E4orf3(-)* Ad-infected HeLa cells collected every 12 h for 48 h and stained with propidium iodide. Bar graphs show mean ± SEM from six independent experiments (n = 6). **(C)** Immunoblot analysis of A549 cells mock-infected or infected with *E4orf3(+)* or *E4orf3(–)* Ad and harvested at the indicated times. Doxorubicin (Dox) and TNFα serve as positive controls. γ-H2AX marks DNA damage; Vinculin is a loading control. (**D, E**) HeLa cells mock-infected or infected with *E4orf3(+)* or *E4orf3(-)* Ad were treated at 4 hpi with (D) the ATM inhibitor KU-60019 (5, 10, 15 µM) or (E) the ATR inhibitor VE-822 (1, 2, 4 µM). At 48 hpi, cell-cycle profiles were analyzed by flow cytometry. Percentages of cells with DNA > 4N are shown as mean ± SEM from three independent experiments. Statistical significance: *p < 0.02, **p < 0.002, ***p < 0.0001. **(F)** HeLa cells mock-infected or infected with E4orf3(+) or E4orf3(–) Ad were treated at 4 hpi with the ATR inhibitor Ceralasertib (0.1, 0.5, 1.0 µM). Abnormal DNA content (DNA > 4N) was quantified at 48 hpi. Data represent mean ± SEM from three independent experiments; *p < 0.02, **p < 0.002, ***p < 0.0001. **(G)** HeLa cells mock-infected or infected with E4orf3(+) or E4orf3(–) Ad were treated at 4 hpi with caffeine (0–12.5 mM). Abnormal DNA content was measured at 48 hpi. Data represent mean ± SEM from three independent experiments; *p < 0.01, **p < 0.001, ***p < 0.0001.

To determine whether ATM and ATR signal through their canonical downstream kinases [1, 2, 16] during infection, we monitored CHK1 and CHK2 phosphorylation in A549 cells (Fig. 1C). Both kinases were phosphorylated in *E4orf3(+)* and *E4orf3(-)* Ad-infections, but only *E4orf3(-)* cells maintained downstream CHK1 and CHK2 signaling (Fig. 1C). This disconnect between ATM/ATR phosphorylation and downstream signaling indicate that checkpoint signaling are rewired in *E4orf3(+)* Ad-infections.

Pharmacological inhibition of ATM (KU-60019) or ATR (VE-822) beginning at 4 hpi revealed that either kinase markedly reduced the fraction of cells with abnormal DNA content (blue bars in Fig. 1D, 1E). In contrast, ATM inhibition caused *E4orf3(-)* Ad-infected cells to accumulate abnormal DNA content (Fig. 1D, red bars), whereas ATR inhibition modestly increased the proportion of cells with abnormal DNA content (Fig. 1E, red bars). These opposing outcomes demonstrate that ATM and ATR have distinct, context-dependent functions, shaped by E4orf3.

To further dissect ATR function, we compared VE-822 with Ceralasertib (AZD6738) [1, 17, 18]. Unlike VE-822, Ceralasertib reduced abnormal DNA content in *E4orf3(-)* Ad infections (Fig. 1F), consistent with reports that these ATR inhibitors exhibit divergent biological activities [19]. As an orthogonal approach, caffeine -an inhibitor of both ATM and ATR [31]- reduced abnormal DNA content in *E4orf3(+)* Ad infections (Fig. 1G) but increased it in *E4orf3(-)* Ad infections (Fig. 1G).

Thus, E4orf3 rewires ATM-ATR signaling, enabling these kinases to promote abnormal DNA content rather than enforcing canonical checkpoint arrest.

### ATM and ATR impose distinct cell-cycle checkpoints governing abnormal DNA content

Because ATM and ATR inhibition produced opposing outcomes depending on E4orf3 status, we next examined how each kinase shapes cell-cycle progression, the process most directly linked to DNA content. In *E4orf3(+)* infections, inhibition of either kinase prevented further accumulation of abnormal DNA content (Fig. 1D-G). ATM inhibition caused a pronounced G2/M accumulation (Fig. 2B, 2C), whereas ATR inhibition blocked progression from G1 to S phase progression (Fig. 2D). These results indicate that ATM and ATR regulate abnormal DNA content through distinct, phase-specific checkpoint functions.

**Figure 2.**
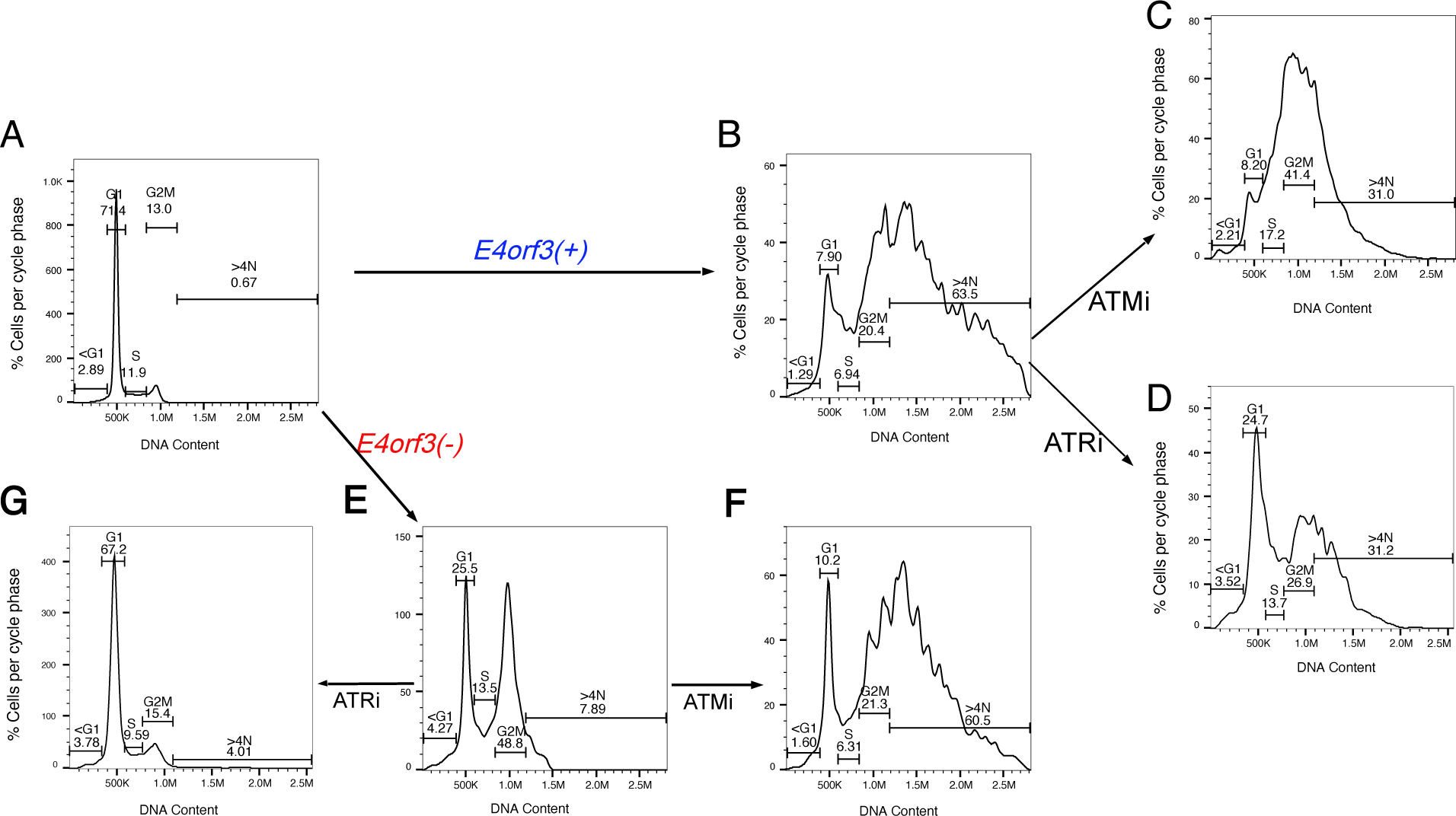
ATM and ATR impose distinct cell-cycle checkpoints governing abnormal DNA content. (**A-G**) HeLa cells were mock-infected or infected with *E4orf3(+)* or *E4orf3(-)* Ad. At 4 hpi, cells were treated with either the ATM inhibitor KU-60019 (15 µM) or the ATR inhibitor VE-822 (4 µM). At 48 hpi, cells were fixed, stained with propidium iodide, and analyzed by flow cytometry. **(A)** Mock-infected cells predominantly reside in G1. **(B)** *E4orf3(+)* Ad-infected cells accumulate high levels of abnormal DNA content (DNA > 4N). **(C)** ATM inhibition reduces abnormal DNA content in *E4orf3(+)* Ad infections and redirects cells into a pronounced G2/M arrest. **(D)** ATR inhibition decreases abnormal DNA content in *E4orf3(+)* Ad infections, with cells accumulating in both G1 and G2/M. **(E)** *E4orf3(-)* Ad-infected cells arrest in G1 and G2/M and show minimal abnormal DNA content. **(F)** ATM inhibition in E4orf3(-) Ad-infected cells induces substantial abnormal DNA content, indicating loss of ATM-dependent G2/M checkpoint control. **(G)** ATR inhibition prevents *E4orf3(-)* cells from progressing through the cell cycle, resulting in G1 accumulation. Representative histograms from three independent experiments are shown.

*E4orf3(-)* infections displayed a bimodal G1 and G2/M distribution (Fig. 1A, 2E). ATM inhibition caused these cells to accumulate a broad range of abnormal DNA content (Fig. 2F), whereas ATR inhibition reduced abnormal DNA content by preventing G1-to-S progression (Fig. 2G). These findings reveal a clear division of labor: ATR safeguards S-phase entry, whereas ATM enforces the G2/M checkpoint. Disrupting ATM-dependent signaling removes a critical barrier to genome replication, enabling cells to accumulate abnormal DNA content.

Thus, ATM and ATR regulate abnormal DNA content by enforcing distinct, phase-specific checkpoints.

### Phosphorylated NEMO links ATM to NF-κB signaling during Ad-infection

The observation that ATM is phosphorylated in line with its activation yet fails to signal to CHK2 in *E4orf3(+)* infections (Fig. 1C) suggests that ATM may redirect its signaling to alternative substrates. Prior work showed that ATM phosphorylation correlates with active NF-κB, which is required for abnormal DNA content in *E4orf3(+)* Ad-infected cells [13]. Because NF-κB activation depends on the IKK complex and its regulatory subunit NEMO, and ATM is known to phosphorylate NEMO [20], we tested whether NEMO mediates NF-κB activation during Ad infection (Fig. 3A)

**Figure 3.**
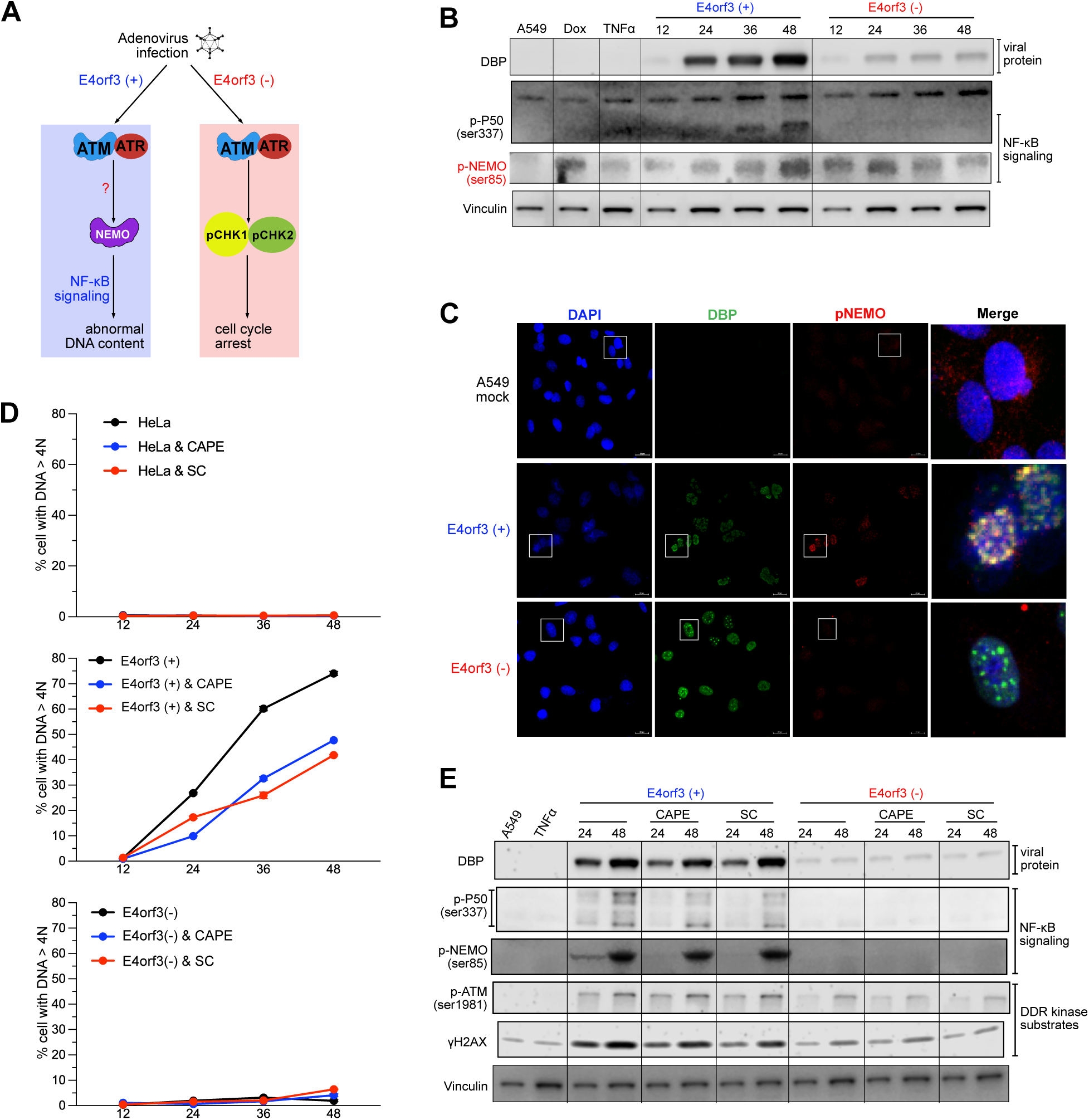
Phosphorylated NEMO links ATM to NF-κB signaling during Ad-infection. **(A)** Schematic illustrating the proposed role of NEMO in mediating NF-κB activation downstream of ATM in *E4orf3(+)* adenovirus (Ad) infection. **(B)** Immunoblot analysis of A549 cells infected with *E4orf3(+)* or *E4orf3(-)* Ad and collected every 12 h up to 48 hpi. Phosphorylation of NEMO and NF-κB p50 increases selectively in *E4orf3(+)* infections. **(C)** Confocal microscopy of A549 cells infected for 48 h with the indicated Ads and stained for Ad DBP (green), phosphorylated NEMO (red), and DAPI (blue). Insets (white boxes) show magnified merged images. Scale bar, 20 µm. **(D)** Flow-cytometric quantification of abnormal DNA content (DNA > 4N) in A549 cells infected with *E4orf3(+)* or *E4orf3(-)* Ad and treated at 4 hpi with or without the NF-κB inhibitors CAPE or Sulfasalazine (SC). Data represent three independent experiments. **(E)** Immunoblot analysis of lysates from the same conditions as in (D), showing effects of NF-κB inhibition on phosphorylation of NEMO, NF-κB p50, and ATM. Results represent three independent experiments.

Western blotting showed that phospho-p50 (serine 337) increased steadily in *E4orf3(+)* Ad infections (Fig. 3B), mirroring the rise in phospho-NEMO and the abnormal DNA content observed in Figs. 1 and 2. In contrast, *E4orf3(–)* infections showed reduced phospho-NEMO and no increase in phospho-p50 (Fig. 3B). Confocal microscopy confirmed that phospho-NEMO localized predominantly in the nuclei of *E4orf3(+)* cells but was nearly undetectable in *E4orf3(-)* and mock-infected control cells (Fig. 3C).

NF-κB inhibition (Caffeic Acid Phenethyl Ester (CAPE) [21] or Sulfasalazine (SC) [22] reduced abnormal DNA content (Fig. 3D) and lowered phospho-p50 levels (Fig. 3E), whereas phospho-NEMO and phospho-ATM levels remained unchanged, indicating that ATM and NEMO act upstream of NF-κB.

These findings identify NEMO phosphorylation as a key event linking ATM activation to NF-κB signaling in *E4orf3(+)* Ad-infected cells and demonstrate that this pathway promotes abnormal DNA content.

### NEMO phosphorylation is required for abnormal DNA content and redirects ATM away from MRN

To test whether NEMO is required for abnormal DNA content (Fig. 4A), we depleted NEMO using siRNAs and confirmed efficient knockdown by immunoblotting (Fig. 4B). NEMO depletion significantly reduced abnormal DNA content in both *E4orf3(+)* and *E4orf3(-)* infections (Fig. 4C), demonstrating that NEMO is essential for this phenotype.

**Figure 4.**
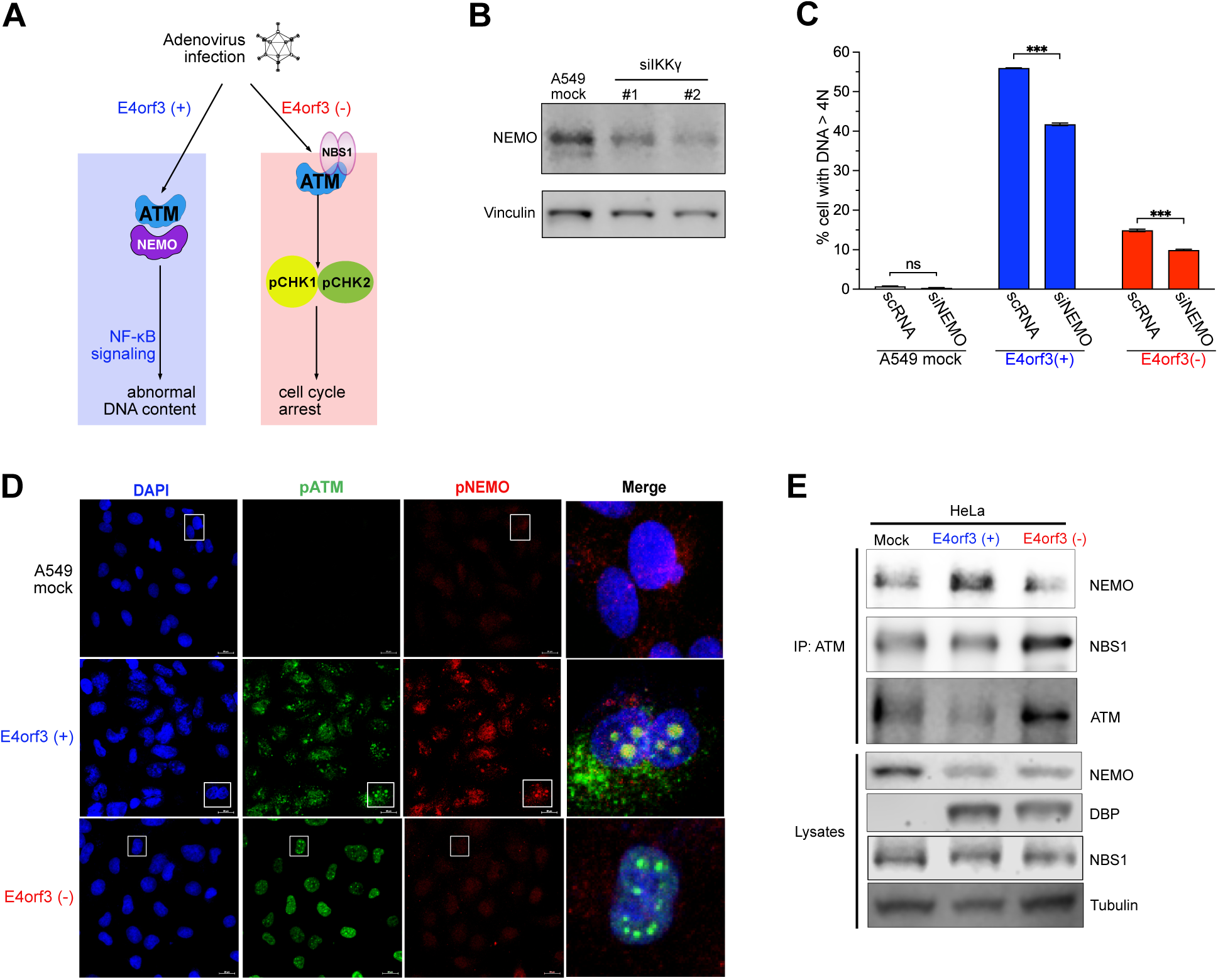
NEMO phosphorylation is required for abnormal DNA content and redirects ATM away from MRN. **(A)** Schematic illustrating the proposed interactions between ATM/ATR and NEMO in *E4orf3(+)* Ad-infected cells. (**B, C**) A549 cells were transfected with control or NEMO-targeting siRNA for 48 hours. **(B)** Immunoblot confirming efficient NEMO depletion. **(C)** Cells were mock-infected or infected with *E4orf3(+)* or *E4orf3(-)* Ads, fixed at 48 hpi, stained with propidium iodide, and analyzed by flow cytometry. NEMO knockdown significantly reduced abnormal DNA content (mean ± SEM, n = 3; ***p < 0.001). **(D)** Immunofluorescence microscopy of A549 cells infected with *E4orf3(+)* or *E4orf3(-)* Ad for 48 h and stained for phosphorylated ATM (green) or phosphorylated NEMO (red), and DAPI (blue). Insets (white boxes) show magnified merged images. Scale bars, 20 μm. **(E)** Co- immunoprecipitation analysis of HeLa cells infected with *E4orf3(+)* or *E4orf3(-)* Ad for 48 h. Lysates were immunoprecipitated and probed for ATM, NEMO, and NBS1, demonstrating preferential ATM-NEMO association in *E4orf3(+)* infections and ATM-NBS1 association in *E4orf3(-)* infections.

Immunofluorescence revealed that phospho-ATM colocalized with phospho-NEMO in *E4orf3(+)* cells (Fig. 4D, row 2), whereas in *E4orf3(-)* cells, phospho-ATM did not colocalize with phospho-NEMO (Fig. 4D, row 3). Co-immunoprecipitation further showed that ATM preferentially associated with NEMO in *E4orf3(+)* infections, while in *E4orf3(-)* infections, it associated with NBS1 (Fig. 4E), a component of the MRN complex [23, 24].

Thus, E4orf3 redirects ATM away from its canonical MRN-dependent functions and toward NEMO, whose phosphorylation is required for NF-κB activation and for abnormal DNA content.

### Disrupting ATM-dependent DDR signaling is sufficient to activate NF-κB and drive abnormal DNA content

Because E4orf3 disrupts the MRN-dependent DDR signaling and redirects ATM toward NEMO, we asked whether blocking ATM’s ability to transmit canonical DDR signals is itself sufficient to activate the NEMO-NF-κB pathway (Fig. 5A). As expected, *E4orf3(+)* Ad infections showed robust phosphorylation of NF-κB p50 (Fig. 5B). Strikingly, ATM inhibition alone induced phosphorylation of NEMO and NF-κB p50 in *E4orf3(-)* Ad-infected cells, in sharp contrast to untreated *E4orf3(-)* infections (Fig. 5B). ATMi inhibition also increased Ad DBP levels, consistent with ATM’s role in restricting proteins required for viral replication [25].

**Figure 5.**
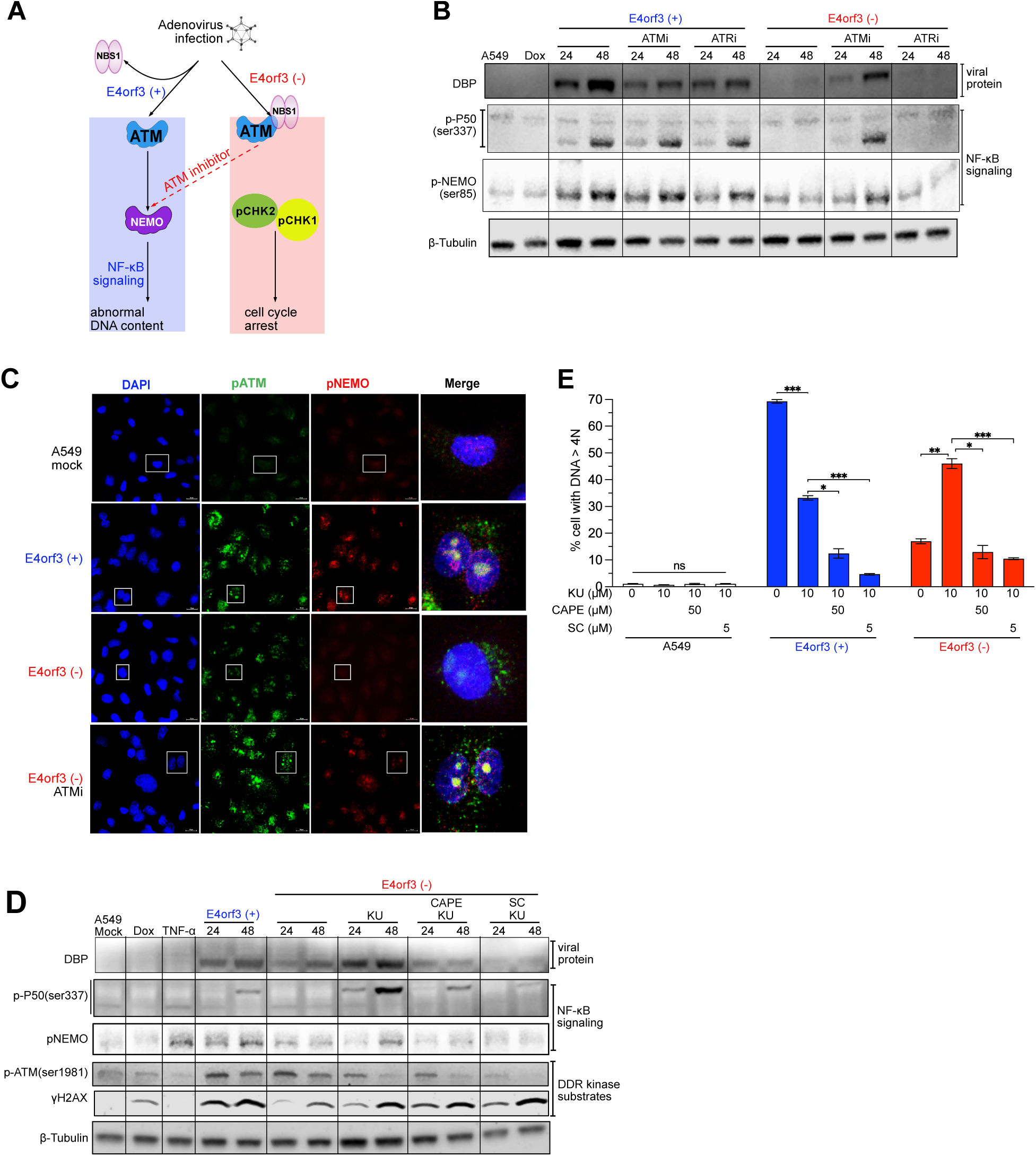
Disrupting ATM-dependent DDR signaling is sufficient to activate NF-κB and drive abnormal DNA content. **(A)** Schematic illustrating how inhibiting ATM in *E4orf3(-)* Ad-infected cells redirects DDR signaling towards NEMO-NF-κB activation. (**B, C**) A549 cells were mock-infected or infected with *E4orf3(+)* or *E4orf3(-)* Ads and treated at 4 hpi with or without ATM or ATR kinase inhibitors. **(B)** Immunoblot analysis of lysates collected at 24 and 48 hpi showing phosphorylation of ATM, NEMO, and NF-κB p50, and accumulation of the Ad DBP. **(C)** Immunofluorescence microscopy of cells fixed at 48 hpi and stained for phosphorylated ATM (green), phosphorylated NEMO (red), and DAPI (blue). Insets (white boxes) show magnified merged images. Scale bar, 20 μm. (**D, E**) A549 cells were mock-infected or infected with *E4orf3(+)* or *E4orf3(-)* Ads and treated 4 hpi with or without ATM inhibitor KU-60019 (KU) and/or the NF-κB inhibitors CAPE or Sulfasalazine (SC). **(D)** Immunoblot analysis of cell lysate collected at 48 hpi showing effects of NF-κB inhibition on phosphorylation of ATM, NEMO, and NF-κB p50. **(E)** Flow-cytometric quantification of abnormal DNA content (DNA > 4N) at 48 hpi. Data represent mean ± SEM from three independent experiments; *p < 0.05, **p < 0.01, ***p < 0.001.

Immunofluorescence analysis showed that phosphorylated NEMO colocalized with phosphorylated ATM in ATMi-treated *E4orf3(-)* Ad-infected cells (Fig. 5C, row 4), mirroring the pattern observed in *E4orf3(+)* Ad-infections (Fig. 5C, row 2). NF-κB inhibition (CAPE or SC) reduced phosphorylation of NF-κB p50, NEMO, and ATM (Fig. 5D) and reversed the ATMi-induced increase in abnormal DNA content (Fig. 5E).

Thus, disabling ATM-dependent DDR signaling is sufficient to activate NEMO-NF-κB and drive abnormal DNA content.

### Disrupting CHK2-dependent DDR signaling is sufficient to activate NF-κB and drive abnormal DNA content

CHK2 is a central effector of MRN- and ATM-dependent DDR signaling [26]. We therefore asked whether disabling CHK2 would mimic ATM inhibition and redirect ATM activity toward NEMO-NF-κB signaling (Fig. 6A).

**Figure 6.**
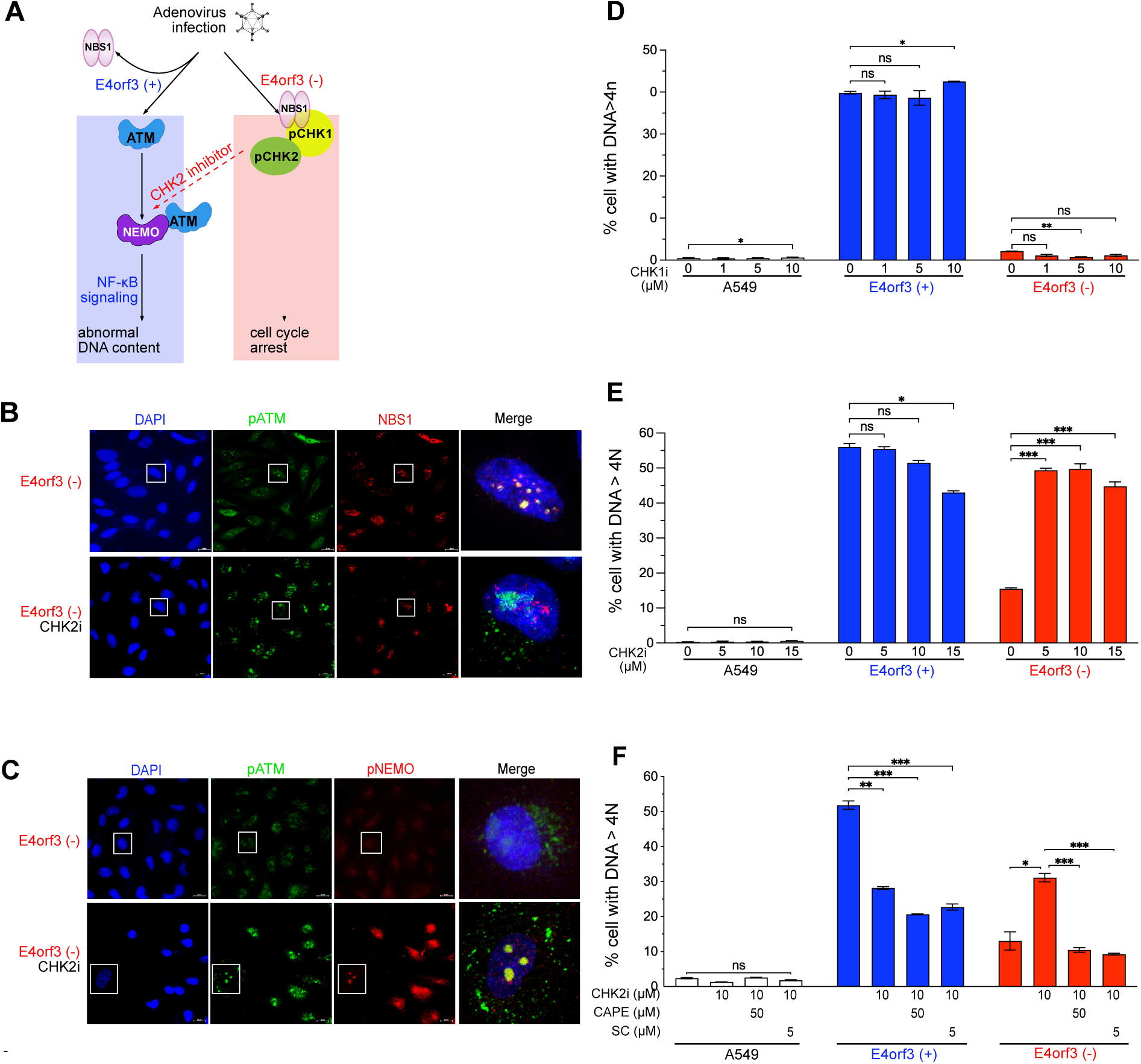
Disrupting CHK2-dependent DDR signaling is sufficient to activate NF-κB and drive abnormal DNA content. **(A)** Schematic illustrating how inhibiting CHK2 in *E4orf3(-)* Ad-infected cells redirects ATM from the MRN complex toward NEMO, thereby activating the NF-κB pathway and promoting abnormal DNA content. **(B)** A549 cells infected with *E4orf3(-)* Ad were treated at 4 hpi with or without CHK2 inhibitors. At 48 hpi, cells were fixed and stained for phosphorylated ATM (green) and NBS1 (red). Insets (white boxes) show magnified merged images. Scale bar, 20 μm. **(C)** A549 cells infected with E4orf3(–) Ad were mock-treated or treated with CHK2 inhibitor at 4 hpi. At 48 hpi, cells were stained for phosphorylated ATM (green), phosphorylated NEMO (red), and DAPI (blue). Insets highlight merged zoomed-in regions. Scale bar, 20 μm. (**D, E**) A549 cells infected with *E4orf3(+)* or *E4orf3(-)* Ad were treated at 4 hpi with increasing concentrations of (D) CHK1 inhibitor or (E) CHK2 inhibitor. At 48 hpi, cells were fixed, stained with propidium iodide, and analyzed by flow cytometry. Percentages of cells with abnormal DNA content (DNA > 4N) are shown as mean ± SEM from three biological replicates; *p < 0.05, **p < 0.01, ***p < 0.001. **(F)** A549 cells infected with *E4orf3(+)* or *E4orf3(-)* Ad were treated at 4 hpi with a CHK2 inhibitor and/or NF-κB inhibitors CAPE or Sulfasalazine (SC). At 48 hpi, abnormal DNA content was quantified by flow cytometry. Data represent the mean ± SEM from three independent experiments.

In untreated *E4orf3(-)* Ad-infections, phosphor-ATM colocalized with NBS1 (Fig. 6B, row 1), consistent with intact MRN-dependent DDR signaling. Strikingly, CHK2 inhibition disrupted ATM-NBS1 colocalization (Fig. 6B, row 2) and instead caused phosphor-ATM to colocalize with phospho-NEMO (Fig. 6C, row 2), indicating that CHK2 is required to maintain ATM’s canonical pathway fidelity.

CHK1 inhibition did not alter abnormal DNA content (Fig. 6D), indicating that CHK1 is not required for this phenotype.

CHK2 inhibition produced opposite outcomes depending on E4orf3 status. In *E4orf3(+)* infections, CHK2 inhibition reduced abnormal DNA content (Fig. 6E, blue bars), similar to ATM inhibition. In contrast, CHK2 inhibition markedly increased abnormal DNA content in *E4orf3(-)* Ad-infected cells (Fig. 6E, red bars). NF-κB inhibition reduced abnormal DNA content in CHK2i-treated *E4orf3(-)* Ad-infected cells (Fig. 6F, red bars), demonstrating that NF-κB activity is essential for CHK2i-treated cells to accumulate abnormal DNA content.

Thus, CHK2 is a critical determinant of ATM pathway fidelity, and its inhibition is sufficient to redirect ATM toward NEMO-NF-κB signaling and promote abnormal DNA content.

### Switching from CHK1/2, to the NEMO-NF-κB pathway induces survival and chemoresistance-associated gene expression

Cells with abnormal DNA content are known to show enhanced viability under stress [27]. Because redirecting ATM/ATR signaling toward NEMO-NF-κB drives abnormal DNA content, we next asked whether this signaling shift also reprograms the transcriptional outputs associated with stress adaptation and chemoresistance.

RNA sequencing of A549 cells infected with *E4orf3(+)* or *E4orf3(-)* Ad and treated with an ATM inhibitor revealed that myeloblastosis proto-oncogene-like 2 (MYBL2) was among the most significantly upregulated genes in both conditions (fold changes = 2.623 and 2.587, respectively; Fig. 7A). MYBL2 is a transcription factor essential for cell cycle progression and survival, and its dysregulation is linked to abnormal DNA content, genomic instability, and poor cancer outcomes [28]. MYBL2 is activated by Cyclin A/E-CDK2 [28], and we previously demonstrated that CDK2 is activated in Ad-infected cells [8].

**Figure 7.**
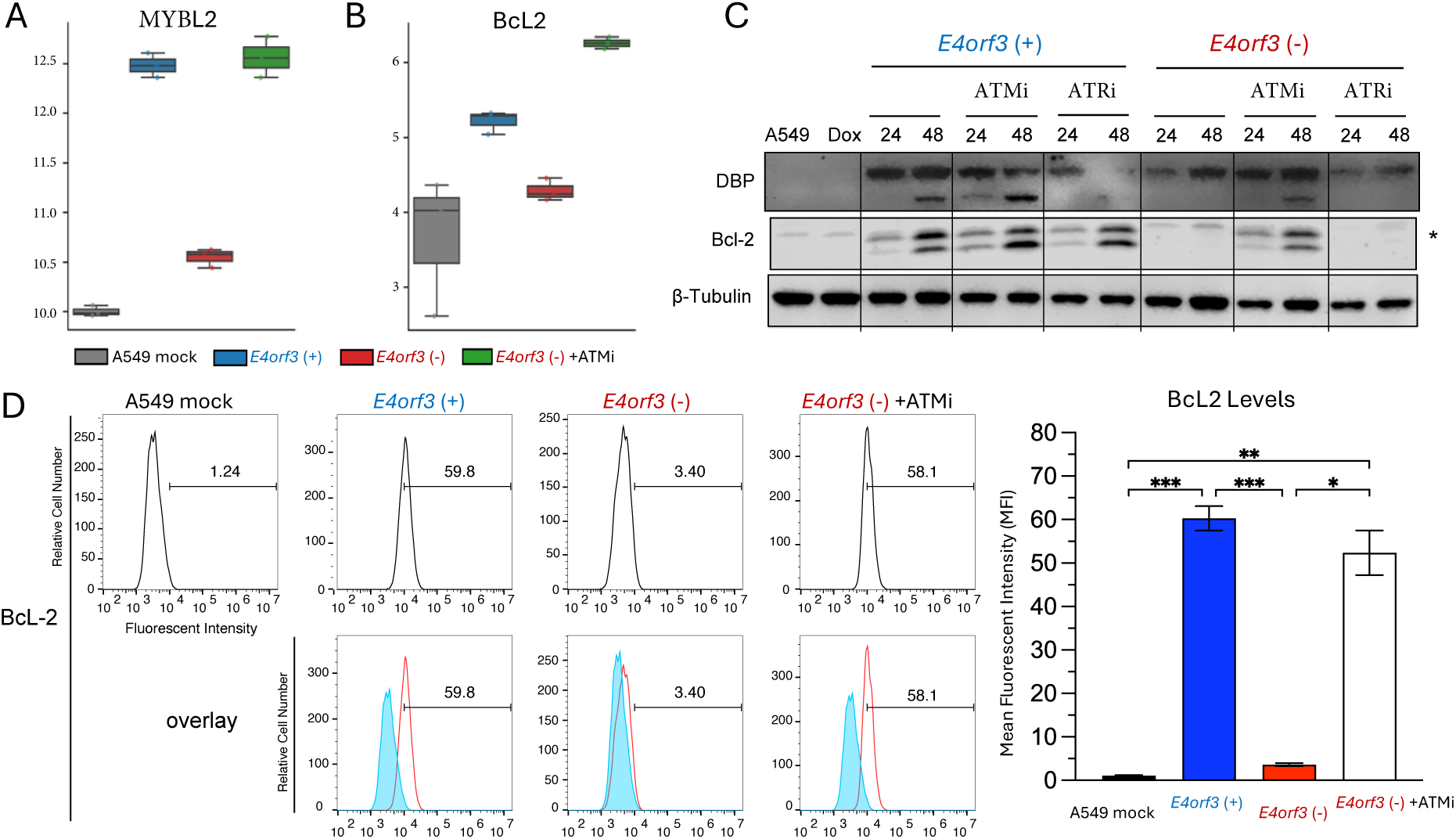
Switching from CHK1/2 to the NEMO-NF-κB pathway induces survival and chemoresistance-associated gene expression. (**A, B**) A549 cells were mock-infected or infected in triplicate with *E4orf3(+)* or *E4orf3(-)* Ad. A subset of *E4orf3(-)* Ad infections received ATM inhibitor (ATMi) at 4 hpi. RNA was isolated, sequenced, normalized for depth, and analyzed using DESeq2. Log₂ fold-changes (mean ± SEM) for differentially expressed genes are shown. **(A)** MYBL2, expression was strongly upregulated in *E4orf3(+)* and ATMi-treated *E4orf3(-)* Ad infections (adjusted p = 3.25 × 10⁻¹⁹² and 6.81 × 10⁻¹⁸⁹, respectively). **(B)** BCL2 expression was similarly elevated in *E4orf3(+)* and ATMi-treated *E4orf3(-)* Ad infections (adjusted p = 7.67 × 10⁻⁶ and 8.98 × 10⁻¹⁶, respectively). **(C)** Immunoblot analysis of A549 cells infected with *E4orf3(+)* or *E4orf3(-)* Ad and treated at 4 hpi with ATMi or ATRi. Lysates collected at 24 and 48 hpi were probed for adenovirus DBP, Bcl-2, and β-tubulin (loading control). **(D)** Flow-cytometric analysis of Bcl-2 protein levels in A549 cells infected with *E4orf3(+)* or ATMi*-*treated *E4orf3(-)* Ad. Top left: representative histograms showing mean fluorescence intensity (MFI). Bottom left: mock overlay on infected samples. Right: quantification of MFI (mean ± SEM, n = 3). Statistical significance: *p < 0.05, **p < 0.005, ***p < 0.001.

MYBL2 promotes survival through multiple mechanisms, including activation of the PI3K/Akt signaling pathway and induction of Bcl-2 [29]. Consistent with this, Bcl-2 RNA levels increased significantly in *E4orf3(+)* Ad-infections (fold change of 1.58) and even more strongly in ATMi-treated *E4orf3(-)* infections (fold change of 2.52) (Fig. 7B). Immunoblotting and flow cytometry confirmed that Bcl-2 protein levels were highest in *E4orf3(+)* and ATMi-treated *E4orf3(-)* Ad-infected cells (Fig. 7C, D).

Thus, redirecting ATM/ATR signaling from CHK1/CHK2 to the NEMO-NF-κB pathway induces transcriptional and protein-level changes characteristic of stress-tolerant, chemoresistant cells, including upregulation of MYBL2, activation of CDK2, and increased Bcl-2 expression.

## DISCUSSION

Adenovirus infection has long been understood to disable canonical DNA-damage response (DDR) signaling by dismantling the MRN complex [11], thereby preventing activation of ATM and ATR [9, 10]. Yet our findings reveal that ATM and ATR are not merely passive bystanders in this context. Instead, Ad, through E4orf3, repurposes these kinases, redirecting their signaling away from the CHK1/CHK2 checkpoint axis and toward the NEMO-NF-κB pathway. This rewiring induces transcriptional programs associated with survival and chemoresistance, revealing a previously unrecognized plasticity in DDR signaling.

A central finding of this study is that ATM activation persists despite MRN disruption, but its downstream signaling is selectively suppressed in *E4orf3(+)* infections. Although ATM and ATR are phosphorylated, their canonical targets, CHK1 and CHK2, are not maintained, revealing a disconnect between kinase activation and checkpoint enforcement. This uncoupling explains the striking divergence in cell-cycle phenotypes: *E4orf3(-)* infections retain a robust G2/M arrest, whereas *E4orf3(+)* infections progress to abnormal DNA content.

Our data show that E4orf3 is sufficient to reprogram ATM/ATR signaling, redirecting these kinases toward NEMO. This is evident in the colocalization of phosphorylated ATM with phosphorylated NEMO, the loss of ATM-NBS1 interactions, and the preferential formation of ATM-NEMO complexes during *E4orf3(+)* infections. In *E4orf3(-)* Ad-infected cells, blocking ATM or CHK2 mimics the effect of E4orf3, demonstrating that the key requirement for pathway switching is not the viral protein itself but the interruption of canonical DDR signaling.

This shift toward NEMO-NF-κB signaling has profound consequences. NF-κB activation is essential for the accumulation of abnormal DNA content, and its inhibition reverses the effects of ATM or CHK2 blockade. These findings position NF-κB as the critical effector of the rewired DDR, transforming what would normally be a checkpoint-enforcing response into a pro-survival, genome-destabilizing program.

RNA-seq analysis revealed strong induction of MYBL2, a transcription factor strongly linked to abnormal DNA content, genomic instability, and poor cancer outcomes [28]. MYBL2 activation, together with persistent CDK2 activity, creates a permissive environment for genome over-replication. Moreover, induction of Bcl-2 at both the RNA and protein levels highlights a shift toward anti-apoptotic signaling. These changes mirror transcriptional programs observed in chemoresistant tumors [30, 31], suggesting that Ad-driven DDR rewiring co-opts pathways typically associated with malignant progression.

Together, these findings support a model in which Ad infection repurposes ATM and ATR from genome guardians into facilitators of viral replication and host-cell survival. By disabling MRN-dependent checkpoint signaling and redirecting ATM/ATR activity toward NEMO-NF-κB, the virus creates a cellular state characterized by abnormal DNA content, enhanced survival, and resistance to stress. This model unifies previously disparate observations-persistent ATM phosphorylation, NF-κB activation, abnormal DNA content, and enhanced viability of infected cells-into a coherent framework for DDR pathway switching.

These insights have important implications for Ad-based therapeutics. E4orf3-containing Ads are being evaluated as oncolytic agents ([32, 33], ClinicalTrials.gov), yet E4orf3-mediated activation of survival pathways may limit their potency. They are also being evaluated for vaccine delivery [34–36]. E4orf3-dependent reprogramming of NF-κB signaling may attenuate NF-κB-driven inflammatory responses that are critical to vaccine efficacy. These considerations underscore the need to account for E4orf3’s signaling effects when designing next-generation Ad vectors.

More broadly, our results reveal a previously unappreciated plasticity in DDR signaling. They demonstrate that ATM and ATR can be reprogrammed not only by viral proteins but also by perturbations to their downstream effectors, such as CHK2. This raises the possibility that similar rewiring events occur in cancer, where DDR components are frequently mutated or suppressed. The parallels between Ad-infected cells and chemoresistant tumor cells—MYBL2 upregulation, CDK2 activation, Bcl-2 induction, and NF-κB dependence—suggest that viruses exploit vulnerabilities that cancer cells also leverage for survival. Understanding this shared logic may illuminate new strategies to counteract therapeutic resistance.

## RESOURCE AVAILABILITY

### Lead Contact

Further information and requests for resources should be directed to

### Materials Availability

This study did not generate new unique reagents.

Materials/Data generated in this study will be shared by the lead contact upon request.

### Data and Code Availability

RNA-seq data will be deposited in a public repository upon acceptance. No custom code was generated for this study.

Microscopy data reported in this paper will be shared by the lead contact upon request. Any additional information required to reanalyze the data reported in this paper is available from the lead contact upon request.

## ACKNOWLEDGMENTS

We thank Dr. David Ornelles (Wake Forest University) for providing the *dl*1520 and *dl*3112 adenoviruses and DBP hybridoma cells. We are grateful to Dr. Marjorie Robert-Guroff for the insightful review of the manuscript. We also thank Dr. Sergei Nekhai (Howard University) for access to the BD FACSVerse, supported by NIH grants 1P50HL118006 and U54MD007597, and Dr. Anna Allen (Howard University) for access to the spinning-disk confocal microscope, funded by a Department of Defense HBCU/MI Equipment/Instrumentation Grant (64684-RT-REP).

## AUTHOR CONTRIBUTIONS

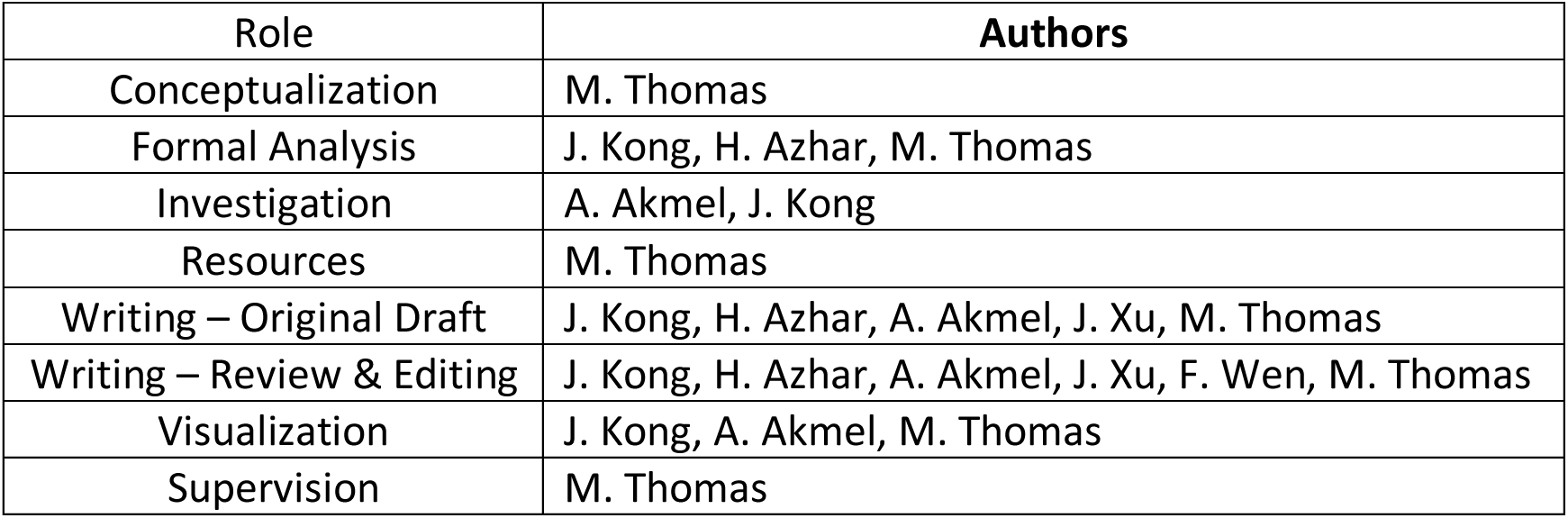
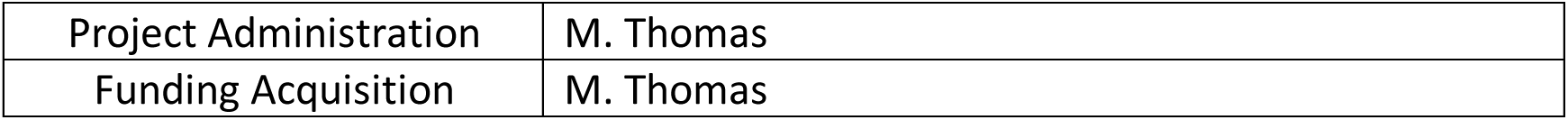

## METHODS

### Key resources table

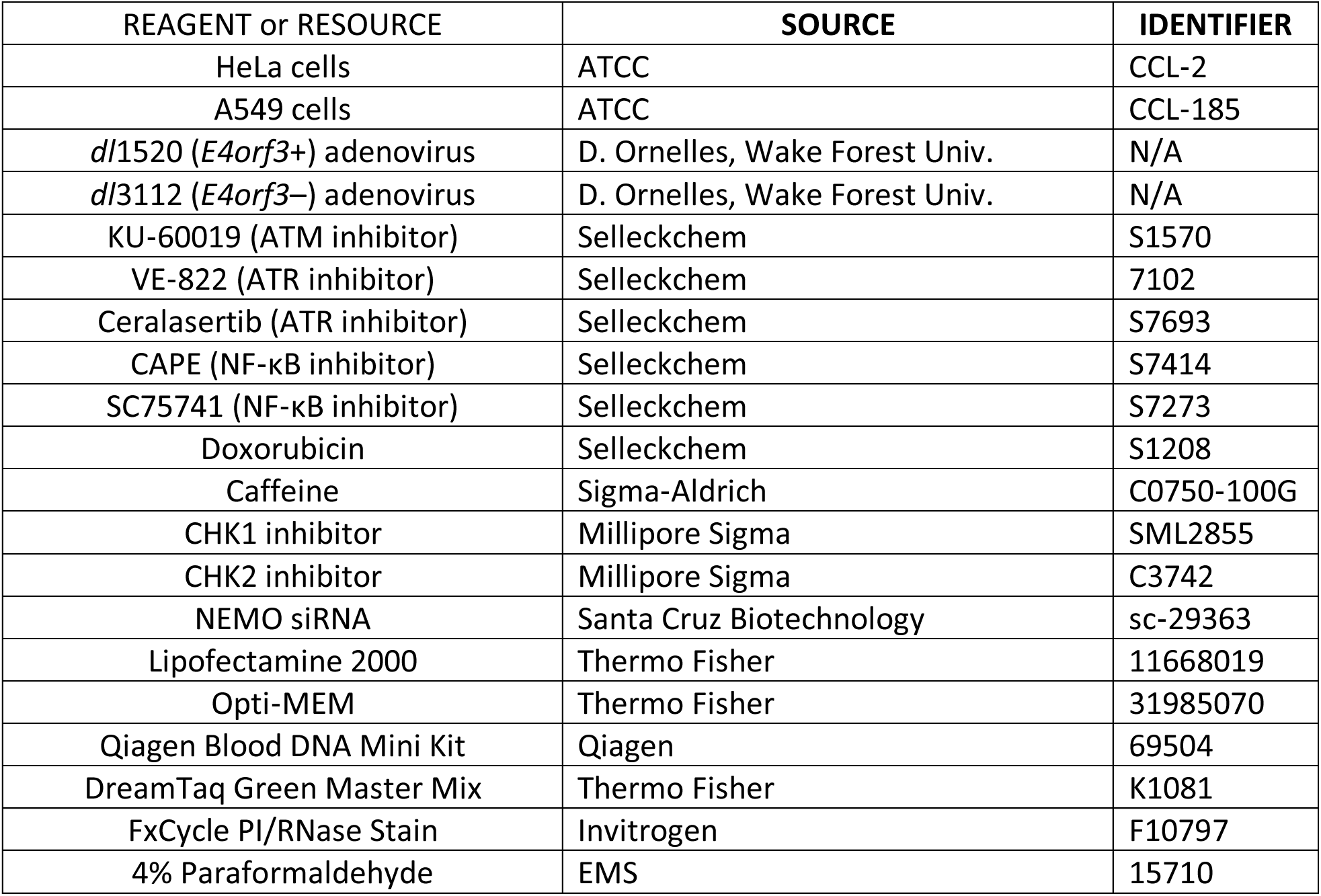
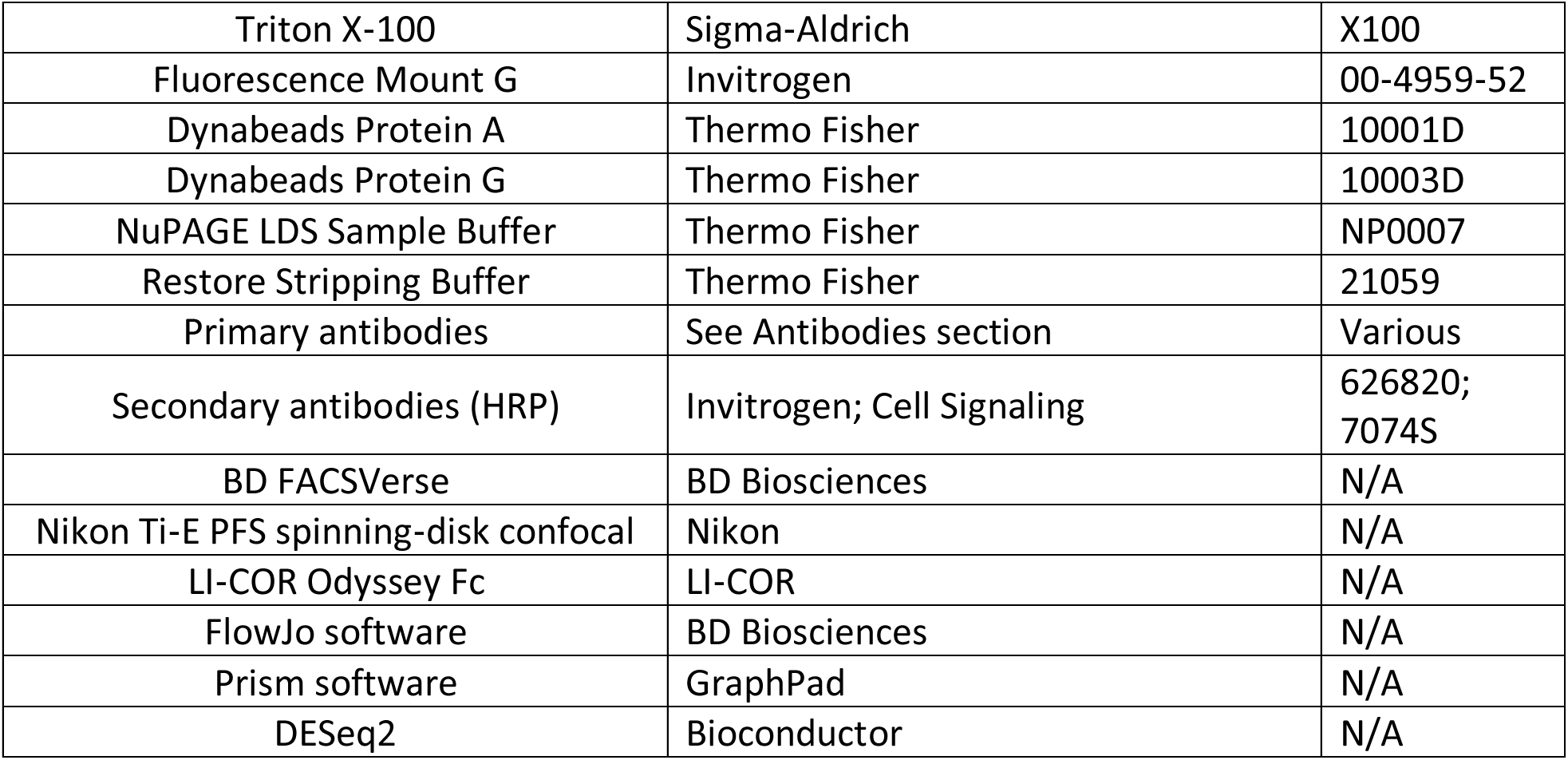

| REAGENT or RESOURCE | SOURCE | IDENTIFIER |
| --- | --- | --- |
| HeLa cells | ATCC | CCL-2 |
| A549 cells | ATCC | CCL-185 |
| *dl1520 (E4orf3+)* adenovirus | D. Ornelles, Wake Forest Univ. | N/A |
| *dl3112 (E4orf3-)* adenovirus | D. Ornelles, Wake Forest Univ. | N/A |
| KU-60019 (ATM inhibitor) | Selleckchem | S1570 |
| VE-822 (ATR inhibitor) | Selleckchem | 7102 |
| Ceralasertib (ATR inhibitor) | Selleckchem | S7693 |
| CAPE (NF-κB inhibitor) | Selleckchem | S7414 |
| SC75741 (NF-κB inhibitor) | Selleckchem | S7273 |
| Doxorubicin | Selleckchem | S1208 |
| Caffeine | Sigma-Aldrich | C0750-100G |
| CHK1 inhibitor | Millipore Sigma | SML2855 |
| CHK2 inhibitor | Millipore Sigma | C3742 |
| NEMO siRNA | Santa Cruz Biotechnology | sc-29363 |
| Lipofectamine 2000 | Thermo Fisher | 11668019 |
| Opti-MEM | Thermo Fisher | 31985070 |
| Qiagen Blood DNA Mini Kit | Qiagen | 69504 |
| DreamTaq Green Master Mix | Thermo Fisher | K1081 |
| FxCycle PI/RNase Stain | Invitrogen | F10797 |
| 4% Paraformaldehyde | EMS | 15710 |
| Triton X-100 | Sigma-Aldrich | X100 |
| Fluorescence Mount G | Invitrogen | 00-4959-52 |
| Dynabeads Protein A | Thermo Fisher | 10001D |
| Dynabeads Protein G | Thermo Fisher | 10003D |
| NuPAGE LDS Sample Buffer | Thermo Fisher | NP0007 |
| Restore Stripping Buffer | Thermo Fisher | 21059 |
| Primary antibodies | See Antibodies section | Various |
| Secondary antibodies (HRP) | Invitrogen; Cell Signaling | 626820;<br>7074S |
| BD FACSVerse | BD Biosciences | N/A |
| Nikon Ti-E PFS spinning-disk confocal | Nikon | N/A |
| LI-COR Odyssey Fc | LI-COR | N/A |
| FlowJo software | BD Biosciences | N/A |
| Prism software | GraphPad | N/A |
| DESeq2 | Bioconductor | N/A |

## EXPERIMENTAL MODEL

### Cell lines

HeLa (ATCC® CCL-2™) and A549 (ATCC® CCL-185) cells were maintained in Dulbecco’s Modified Eagle’s Medium supplemented with 10% fetal bovine serum (FBS), 100 units/ml penicillin, and 100 mg/ml streptomycin at 37°C in a humidified atmosphere with 5% CO_2_. Cells were passaged two or three times weekly at approximately 1:10 dilution.

### Ethics Statement

All experiments were approved by the Howard University Institutional Biosafety Committee (Approval # IBC-2023-0022, IBC-2024-0067).

## METHOD DETAILS

### Viruses

The early region 1B 55K (E1B55K)-deleted adenovirus dl1520, also known as Onyx-015, Lontucirev, and CI-1042 [44], and the E1B55K- and early region 4 (E4) open reading frame 3 (E4orf3)-deleted Ad5 dl3112 [45] have been previously described. In Table 1 and throughout the manuscript, these viruses are designated *E4orf3(+)* and *E4orf3(−),* respectively, to highlight their major differences. All viral stocks were purified, and titers were determined by plaque assay.

**Table 1:** List of viruses.

| Sample | Description | Abnormal DNA Content |
| --- | --- | --- |
| Mock | Cells with no virus | No |
| *E4orf3(+)* | *E1B55K-deleted*, (dl1520, Onyx-015, Lontucirev, CI-1042) | Yes |
| *E4orf3(-)* | *E1B55K, E4orf3-deleted* (dl3112) | No |
| *E4orf3(-)* +ATMi | dl3112 + ATM inhibitor | Yes |

### Infection

Cells were seeded 24 h before infection at 5 × 10⁵ cells/well (6-well) or 2.5 × 10⁵ cells/well (12-well). Infections were performed at an MOI of 30 PFU/cell. The virus was adsorbed for 1 h at 37 °C with gentle rocking, after which the inoculum was replaced with complete medium. Cells were incubated for 24-48 h post-infection (hpi) depending on the experiment.

### Pharmacological Inhibitors

Inhibitors used included: KU-60019 (ATM; Selleck S1570), VE-822 (ATR; Selleck 7102), ceralasertib (ATR; Selleck S7693), CAPE (NF-κB DNA-binding; Selleck S7414), SC75741 (NF-κB nuclear translocation; Selleck S7273), doxorubicin (Selleck S1208), caffeine (Sigma C0750-100G), CHK1 inhibitor (Sigma SML2855), and CHK2 inhibitor (Sigma C3742). Inhibitors were added at indicated time points to interrogate ATM/ATR pathway switching.

### Silencing RNA (siRNA)

Cells were seeded at 2 × 10⁵ cells/well (12-well) in antibiotic-free medium 24 h before transfection. NEMO-targeting siRNA (Santa Cruz sc-29363) was transfected using Lipofectamine 2000 (Thermo Fisher 11668019) in Opti-MEM (Thermo Fisher 31985070) for 4 h, after which medium was replaced with complete antibiotic-free medium. Cells were mock-infected or Ad-infected the following day and harvested at 48 hpi for flow cytometry or immunoblotting to assess NEMO-dependent NF-κB signaling.

### DNA extraction and PCR

Genomic DNA was isolated using the Qiagen Blood DNA Mini Kit. DNA was amplified using DreamTaq Green Master Mix with adenovirus fiber primers in a 25 µL reaction. Each run included a mock-DNA negative control and a mock-DNA sample spiked with Ad DNA as a positive control. Products were resolved on 1% agarose, stained, and imaged on a LI-COR system.

### Flow Cytometry

#### Cell-cycle analysis

Cells were seeded at 2 × 10⁵ cells/well (12-well), infected as described, and harvested at the indicated time points. Cells were fixed overnight in 70% ethanol at -20 °C, washed, and stained with FxCycle PI/RNase solution for 30 min. DNA content (10,000 events per sample) was measured on a BD FACSVerse and analyzed with FlowJo.

#### Intracellular staining

Cells were infected as above, collected at defined time points, fixed in PFA, permeabilized, and stained with fluorescent IgG or anti-Bcl-2 antibodies to quantify intracellular protein levels.

### Immunoblotting

Cells were lysed in 1× SDS sample buffer containing protease/phosphatase inhibitors and 5% β-mercaptoethanol. Equal protein amounts were resolved on 4-20% Tris-Glycine gels and transferred to nitrocellulose, blocked in PBS/0.02% Tween-20 with 20% milk, and incubated with primary antibodies overnight at 4 °C. After washing, membranes were incubated with HRP-conjugated secondary antibodies and developed using a 1:1 luminol/enhancer substrate. Images were acquired on a LI-COR Odyssey Fc. Blots were stripped and reprobed as needed. Ad DBP served as an infection control; β-tubulin or vinculin served as a loading control.

### Antibodies

Primary antibodies: ATM (Millipore Sigma A1106, 1:1000), phospho-ATM Ser1981 (Invitrogen MAB-3806, 1:1000), phospho-ATR Ser1989 (Invitrogen MA5-3626, 1:1000), Bcl-2 (Cell Signaling 15071S, 1:1000), β-tubulin (Invitrogen PA1-16947, 1:5000), phospho-CHK1 Ser345 (Cell Signaling 13303, 1:1000), phospho-CHK2 Thr68 (Invitrogen PA5-17818, 1:1000), phospho-H2AX Ser139 (BioLegend 613402, 1:1000), NBS1 (BD 611870, 1:1000), NEMO (ProSci 2335, 1:1000), phospho-NEMO Ser85 (LSBio LS-C353075, 1:1000), phospho-p50 Ser337 (Santa Cruz SC-271908, 1:1000), and vinculin (Santa Cruz SC-25336, 1:1000). HRP-conjugated secondary antibodies: goat anti-mouse (Invitrogen 626820, 1:5000) and goat anti-rabbit (Cell Signaling 7074S, 1:5000).

### Immunofluorescence

Cells were seeded at 2.5 × 10⁵ cells/well, infected, fixed in 4% PFA, permeabilized with 0.2% Triton X-100, blocked in 10% BSA, and incubated with primary antibodies for 1 h. Alexa Fluor 488- or 561-conjugated secondary antibodies (1:2000) were incubated for 1 h. Samples were mounted with Fluorescence Mount G and imaged on a Nikon Ti-E PFS spinning-disk confocal using a 100× 1.4 NA objective and an Andor iXon 897 EMCCD camera. Images were processed in NIS-Elements software.

### Immunoprecipitation

Cells were mock-infected or Ad-infected in 6-well plates and harvested at 48 hpi. Lysates were prepared in lysis buffer, clarified, and incubated overnight at 4 °C with Dynabeads Protein A/G pre-bound to 5 µg primary antibody. Beads were washed, and bound proteins were eluted in NuPAGE LDS sample buffer with reducing agent at 70 °C for 10 min and analyzed by immunoblotting.

### RNA Sequencing

A549 cells were infected in triplicate at an MOI of 30, collected, snap-frozen, and submitted to Azenta Life Sciences for RNA extraction and sequencing. Read counts were normalized for sequencing depth. Sample similarity was assessed using Euclidean distance. Differential expression was analyzed using DESeq2 with Wald test-derived p-values. Genes with adjusted p < 0.05 and |log₂FC| > 1 were considered differentially expressed, enabling identification of survival and chemoresistance programs.

## QUANTIFICATION AND STATISTICAL ANALYSIS

Data are presented as mean ± SEM. Statistical analyses were performed in Prism. Two-way ANOVA with Tukey’s post-hoc test was used for comparisons involving multiple groups. Significance was defined as p < 0.05.

## SUPPLEMENTAL INFORMATION

### Supplementary Data

**Figure S1:**
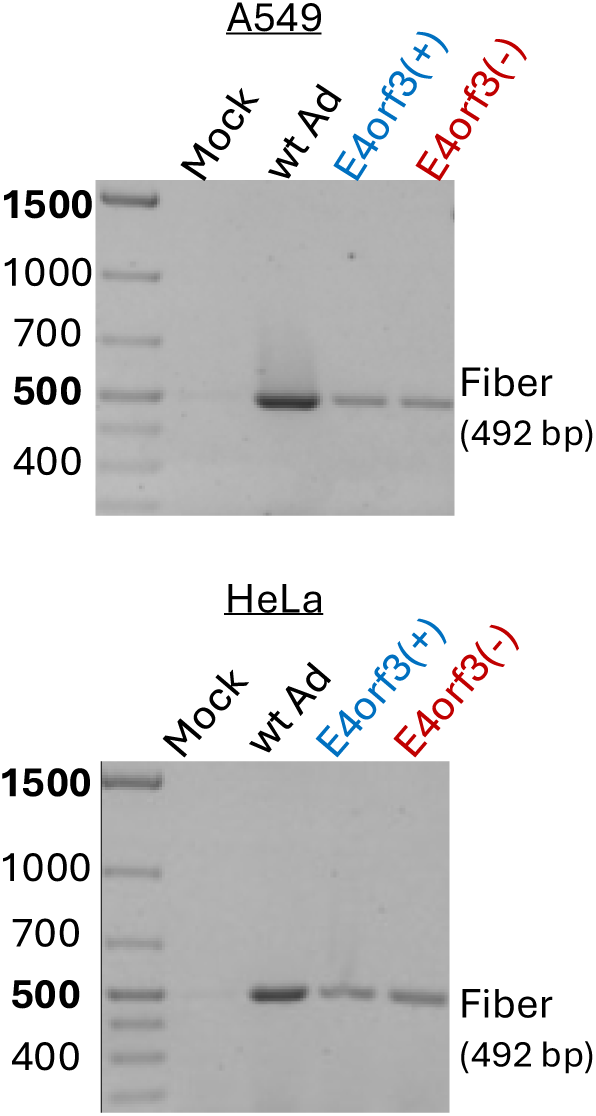
Ad DNA levels are equivalent in *E4orf3(+)* and *E4orf3(-)* infections early after entry. DNA was isolated from A549 and HeLa cells 2 h post-infection (hpi) with *E4orf3(+)* or *E4orf3(-)* Ad, and viral DNA abundance was measured by PCR using Ad fiber–specific primers (Forward: CACCCCTCACAGTTACCTCAGAAGCCC; Reverse: GTCTGTTTTGAGAATCAATCCTTAGTCCTC). Mock-infected samples served as negative controls; wild-type Ad DNA-spiked mock samples served as positive controls. Despite differences in Ad DNA-binding protein (DBP) accumulation observed at later time points (Figs 1D, 3B, 3D), E4orf3(+) and E4orf3(-) infections exhibited comparable viral DNA levels at 2 hpi, indicating equivalent early viral entry and genome delivery.

